# SARS-CoV-2 binding and neutralizing antibody levels after vaccination with Ad26.COV2.S predict durable protection in rhesus macaques

**DOI:** 10.1101/2021.01.30.428921

**Authors:** Ramon Roozendaal, Laura Solforosi, Daniel Stieh, Jan Serroyen, Roel Straetemans, Frank Wegmann, Sietske K. Rosendahl Huber, Joan E. M. van der Lubbe, Jenny Hendriks, Mathieu le Gars, Liesbeth Dekking, Dominika N. Czapska-Casey, Nuria Guimera, Sarah Janssen, Sarah Tete, Abishek Chandrashekar, Noe Mercado, Jingyou Yu, Wouter Koudstaal, Jerry Sadoff, Dan H. Barouch, Hanneke Schuitemaker, Roland Zahn

## Abstract

The first COVID-19 vaccines have recently gained authorization for emergency use.^1,2^ At this moment, limited knowledge on duration of immunity and efficacy of these vaccines is available. Data on other coronaviruses after natural infection suggest that immunity to SARS-CoV-2 might be short lived,^3,4^ and preliminary evidence indicates waning antibody titers following SARS-CoV-2 infection.^5^ Here we model the relationship between immunogenicity and protective efficacy of a series of Ad26 vectors encoding stabilized variants of the SARS-CoV-2 Spike (S) protein in rhesus macaques^6,7,8^ and validate the analyses by challenging macaques 6 months after immunization with the Ad26.COV2.S vaccine candidate that has been selected for clinical development. We find that Ad26.COV2.S confers durable protection against replication of SARS-CoV-2 in the lungs that is predicted by the levels of S-binding and neutralizing antibodies. These results suggest that Ad26.COV2.S could confer durable protection in humans and that immunological correlates of protection may enable the prediction of durability of protection.

---

We previously characterized the immunogenicity and protective efficacy of various Ad26-based vaccine candidates in a rhesus macaque (*Macaca mulatta*) challenge model of SARS-CoV-2^6,7,8^ that resulted in selection of Ad26.COV2.S as the lead vaccine candidate.^9^ To get an early understanding on the potential durability of protection mediated by Ad26.COV2.S, we explored whether immunological markers can be used to predict duration of protection against SARS-CoV-2 in macaques. The absence of detectable viral load was used as a measure of protection. Immunogenicity and efficacy were analyzed across all tested Ad26-based vaccine candidates^6^. This assumes that the immune responses induced by the lead candidate and the prototypes are qualitatively similar, which will be considered in the analysis by comparing the dataset of all Ad26-based vaccine candidates combined with that of Ad26.COV2.S alone. SARS-CoV-2 Spike (S) protein immunogenicity as assessed by psVNA and ELISpot at 4 weeks after vaccination were already reported.^6^ Here, we report additional S-ELISA binding antibody data from these studies (Fig. S1), in an assay homologous to the one utilized for human immunogenicity assessment,^9^ to support the current correlate analysis. The estimated mean probability of protection against detectable viral load (sub-genomic mRNA (sgRNA)), together with a 95% confidence interval (CI), as a function of the level of different immune responses, was obtained using logistic regression, similarly as previously used for anthrax^10^ and Ebola virus disease^11^ vaccines (see statistical methods for more details).

Higher levels of SARS-CoV-2 neutralizing antibodies were associated with an increased protection against viral replication in the lung (Fig. 1a) and the nose (Fig. 1e), both for all vaccine candidates (black), and for Ad26.COV2.S alone (green). The slope of the logistic model for Ad26.COV2.S alone did not have a slope significantly different from 0 (p=0.071), in contrast to the combined model based on all Ad26-based vaccine candidates (p=0.0065), which is likely due to the limited number of animals vaccinated with Ad26.COV2.S that have breakthrough infection. The model based on Ad26.COV2.S had similar high discriminatory capacity in both the lung and the nose, as observed by the area under the ROC curve (AUC = 0.896 and 0.931 respectively).

**Figure 1.**
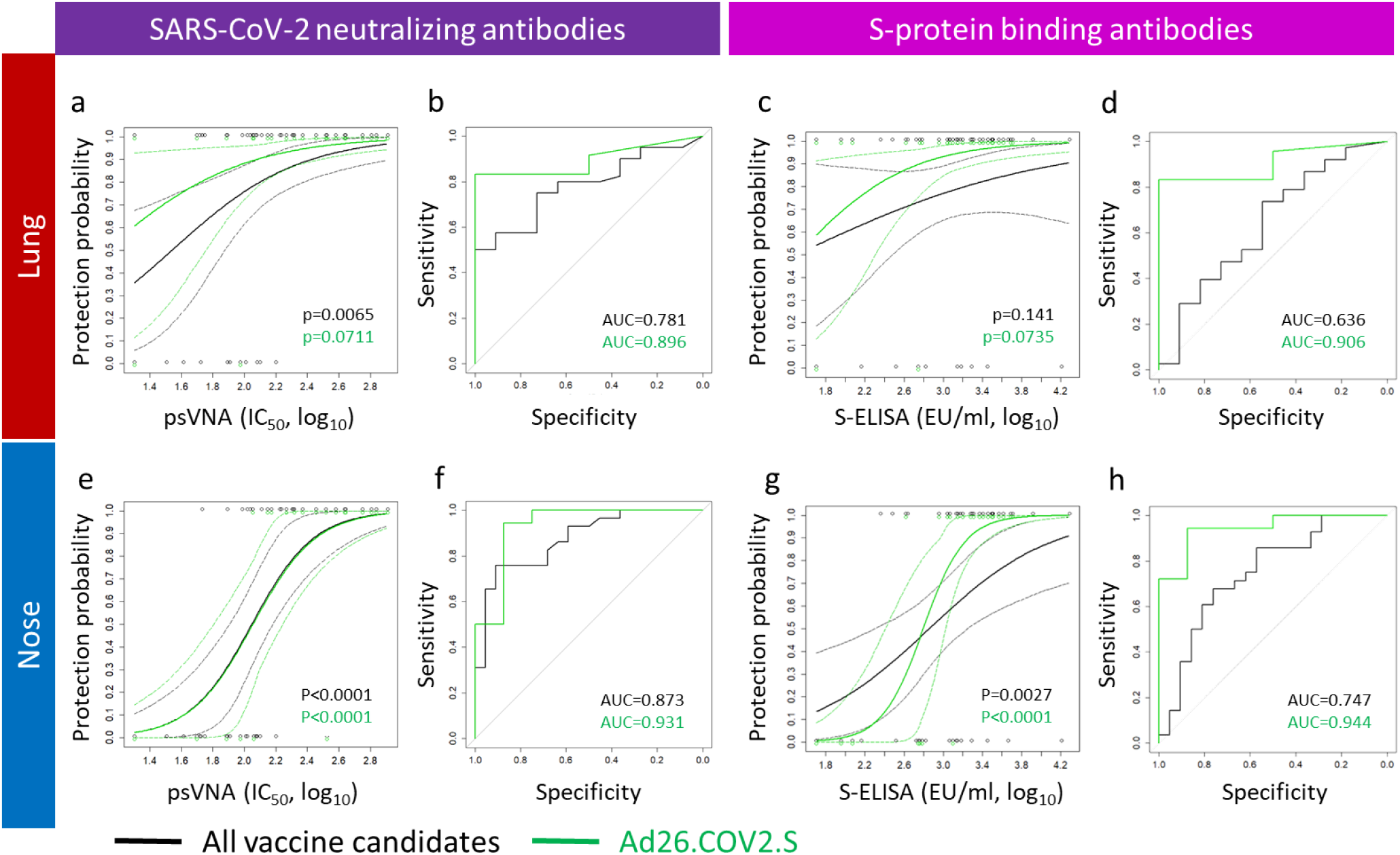
Ad26.COV2.S and prototype vaccines induced levels of SARS-CoV-2 binding and neutralizing antibodies correlate with protection against viral replication in the lung and the nose. (**a, e**) Logistic models of the correlation between the level of SARS-CoV-2 neutralizing antibodies (psVNA at 4 weeks after vaccination, IC_50_, log_10_) and protection against viral load in lung (BAL)(**a**)) and nose (swabs,(**e**)), based on the dataset of all Ad26-based vaccine candidates combined (black line) and Ad26.COV2.S alone (green line). 95% confidence intervals are represented by dashed lines in the same color. Individual datapoints (y=0: detectable viral load; y=1 undetectable viral load) are represented by open circles in the same color. (**b**) ROC curves for the logistic models presented in panel **a**. (**f**) ROC curves for the logistic models presented in panel **e**. Area under the ROC curve (AUC) is indicated and represents a measure of the sensitivity and specificity of the logistic model. (**c, g**) Logistic models of the correlation between the level of S-protein binding antibodies (ELISA at 4 weeks after vaccination, EU/ml, log_10_) and protection against viral load in lung (BAL(**c**)) and nose (swabs,(**g**)), based on the dataset of all vaccine candidates combined (black line) and Ad26.COV2.S alone (green line). 95% confidence intervals are represented by dashed lines in the same color. Individual datapoints (y=0: detectable viral load; y=1 undetectable viral load) are represented by open circles in the same color. (**d**) ROC curves for the logistic models presented in panel **c**. AUC under the ROC is indicated. (**h**) ROC curves for the logistic models presented in panel **g**. AUC under the ROC is indicated.

Similar to neutralizing antibody levels, higher levels of S-protein binding antibodies were associated with an increased protection against viral replication in the lung (Fig. 1c) and nose (Fig. 1g). Only the logistic models based on the nasal viral load data have significant slopes for both all vaccine candidates combined (black, p=0.0027) and Ad26.COV2.S alone (p<0.0001), while the logistic models based on lung viral load data had not, due to the high frequency of complete protection in the lung for Ad26.COV2.S. In addition, the logistic models based on S-ELISA appear to be more sensitive to candidate specific antigenicity than the models based on psVNA, as observed by the more pronounced differences between models based on all vaccine candidates combined compared to Ad26.COV2.S alone (compare Fig. 1e and 1g). Models based on Ad26.COV2.S alone have a higher ROC AUC, both with regards to protection probability in the lung (Fig. 1d, AUC = 0.906 for Ad26.COV2.S vs 0.636 for all vaccine candidates) and the nose (Fig. 1h, AUC = 0.944 for Ad26.COV2.S vs 0.747 for all vaccine candidates). Thus, the logistic models based on Ad26.COV2.S alone had a higher discriminatory capacity than the models based on all vaccine candidates combined in all instances (Fig. 1b, 1d, 1f, and 1h). The high AUC (up to 0.944) indicates that these models should have substantial discriminatory capacity for predicting protection against viral replication. Remarkably, both the binding and neutralizing antibody logistic models based on Ad26.COV2.S predict 60% protection based on immune response levels just above the limit of detection.

Cellular immune responses measured by IFN-γ ELISpot poorly correlated with protection in both lung and nose (Fig. S2A left, and C, left respectively) when analyzed across all vaccine candidates. The dataset based on Ad26.COV2.S alone shows that higher cellular responses are associated with decreased viral load in both the lung and the nose (green curves). However, even for Ad26.COV2.S alone, these models do not have a significant slope, and only the model for protection in the lung has a good discriminatory capacity (AUC=0.875). We therefore focused on the logistic models for binding and neutralizing antibodies as these have a higher discriminatory capacity and are consistent between the dataset across all vaccine candidates and the dataset for Ad26.COV2.S alone.

We subsequently assessed whether vaccination with Ad26.COV2.S provides protection in macaque at 6 months after the first vaccination, and whether the degree of protection could have been anticipated based on the derived correlates of protection. Groups of 7 macaques were vaccinated with either a one-dose (5×10^10^ vp or 1×10^11^ vp) or two-dose regimen (5×10^10^ vp/dose) of Ad26.COV2.S with either a 4-week or an 8-week interval between doses (Fig. 2A)^7^. Twenty-six (26) weeks after the first vaccination animals were challenged with 1×10^5^ TCID50 SARS-CoV-2 via the intranasal and intratracheal routes.^12,13^ Viral loads in BAL and nasal swabs were assessed by reverse transcription PCR (RT–PCR) specific for sgRNA, which predominantly detects replicating virus.^12,14^ All controls had detectable virus in the upper and lower airways at comparable peak levels as in the previous studies.^6,8^ Ad26.COV2.S vaccinated macaques were robustly protected against viral replication in the lung 6 months after first vaccination (Fig. 2b) and the limited lung breakthrough infection in 3 out of 28 macaques was unrelated to either a single or two dose vaccine regimen being applied (Fig. S3). In contrast, most vaccinated animals (24 out of 28) had detectable virus in the nose (Fig. 2c), though duration of virus shedding was significantly lower in the one-dose 1×10^11^ vp group and in the 8-week 2-dose regimen group (Fig. S4, p=0.012) while the latter also showed lower peak viral load (p=0.038) than controls.

**Figure 2.**
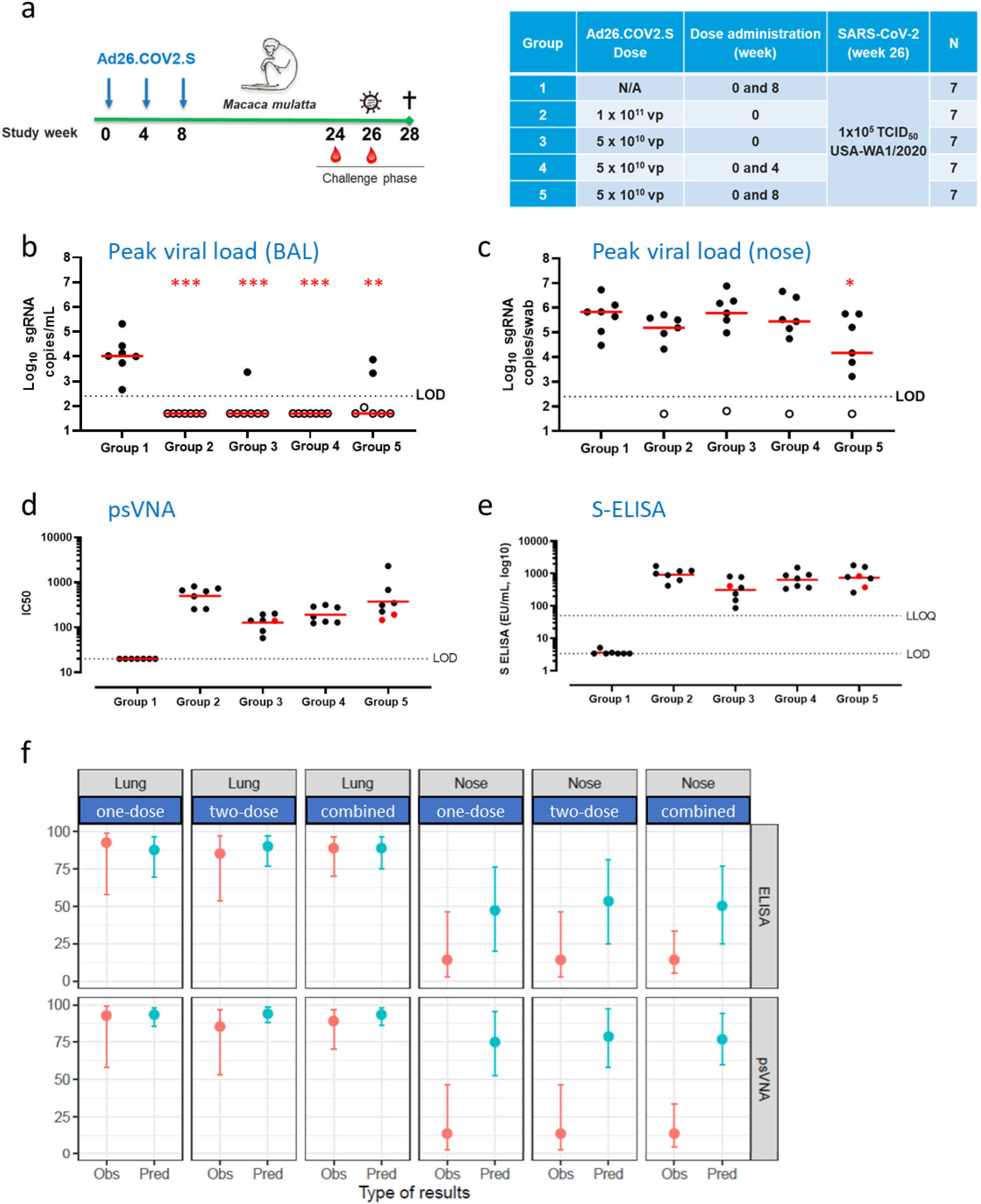
Durable protection against SARS-CoV-2 in the lower airways after vaccination with Ad26.COV2.S is predicted by binding and neutralizing antibody levels. **(a)** Schematic representation of study Groups of 7 monkeys received either a one-dose (5×10^10^ vp or 1×10^11^ vp) or two-dose regimen (5×10^10^ vp/dose) of Ad26.COV2.S with either a 4-week or an 8-week interval between doses. A group of 7 controls was included in the study, of which 4 received 2-doses of saline, and 3 were unvaccinated.

Twenty-six (26) weeks after the first vaccination animals were challenged with 1×10^5^ TCID50 SARS-CoV-2 via the intranasal and intratracheal routes. Peak viral loads in BAL (**b**) and nasal swabs (**c**) were assessed by reverse transcription PCR (RT-PCR) specific for sub-genomic mRNA (sgRNA, copies/mL, log_10_). Assay LOD is indicated by a dashed line. Red line represents group median (**d**) SARS-CoV-2 neutralizing antibodies (psVNA, IC_50_, log_10_) were determined at 24 weeks post vaccination (9-days prior to challenge). Assay LOD is shown as a dashed line. Red line represents group geometric mean titers (GMT) (**e**) S-protein binding antibody levels (S-ELISA, EU/ml, log_10_) were determined at 26 weeks post-vaccination (1 day prior to challenge). Assay LOD and lower limit of quantification (LLOQ) are shown as dashed lines. Red line represents group GMT (**f**) Comparison of the observed (Obs) protection proportion 6 months after vaccination with Ad26.COV2.S with the predicted (Pred) protection probability based on pre-challenge binding (S-ELISA) and neutralizing antibody (psVNA) levels and correlate of protection models (logistic models) constructed based on immunogenicity data of Ad26.COV2.S obtained 4 weeks after vaccination. Asterisks indicate significant difference in peak viral load (*p<0.05, ** p<0.01, *** p<0.001).

Immune response kinetics of this cohort up to 14 weeks after the first immunization are described elsewhere.^7^ We here only consider the levels of SARS-CoV-2 neutralizing antibodies (Fig. 2D) and S-protein binding antibodies (Fig. 2E), measured just prior to challenge, using the same assays used in construction of the logistic models. We focus on logistic models based on Ad26.COV2.S, as this is the same vaccine candidate as used in the durability study.

Data from the durability study were grouped by one-dose regimes (one-dose, n=14), two-dose regimens (two-dose, n=14), or analyzed together (combined, n=28). For each group, the observed proportion of protection (Obs), together with its 95% CI, after challenge (blue dot and interval in Fig. 2f) are compared to the predicted (Pred) mean probability of protection and a bootstrap derived 95% CI, based on their pre-challenge immunogenicity results and the previously developed logistic regression models (red dot and interval in Fig. 2f). For the one-dose regimens, there is excellent agreement between the predicted and the observed protection probabilities in the lung, both when based on S-protein binding antibodies (Pred (88% (69.4-96.4%) vs Obs ((92.9% (58-99.2%)) as well as based on neutralizing antibodies (Pred (93.5% (85.4-97.9%)) vs Obs (92.9% (58-99.2%)).

The observed protection proportion for the two-dose regimens is slightly lower, though still well within the 95% CI of the predicted probability when based on S-protein binding antibodies (Obs 85.7% vs Pred (90.5 (76.7-96.8%)). Overall, the predictions based on binding and neutralizing antibodies show a remarkable correspondence to the observed protection proportion in the lung, indicating that the potential correlates of protection identified early after vaccination can be used to predict durable protection against infection of the lower airways in rhesus macaques.

The observed protection probability in the nose is substantially lower than predicted based on either pre-challenge binding or neutralizing antibody levels in the circulation. While the predicted protection probability was approximately 50% based on S-protein binding antibody levels and 75% based on neutralizing antibody levels, nose protection was only observed in 4/28 animals (14.3%). This shows that systemic binding and neutralizing antibody levels likely are associated with a distinct mechanistic correlate of protection in the nose early after vaccination, rather than being a mechanistic correlate of protection themselves.^15,16^

Predictions for the probability of protection, using logistic models based on data of all vaccine candidates combined largely confirm predictions based on Ad26.COV2.S (Fig. S5), demonstrating the robustness of predicting durable protection in the lower airways based on binding and neutralizing antibody levels. In fact, the model based on binding antibody responses of all vaccine candidates combined under-predicts the observed protection in the lung, reinforcing selection of the Ad26.COV2.S candidate for clinical evaluation of protective efficacy.

In conclusion, one or two doses of Ad26.COV2.S confer a high degree of protection against SARS-CoV-2 replication in the lower airways of macaques for at least 6 months. It is conceivable that this would translate into durable protection against COVID-19 in vaccinated humans, where protection may be even more durable as infection with SARS-CoV-2 is likely to trigger an anamnestic response.^17^ Due to the incubation time between infection and development of symptoms, humans may indeed benefit from an anamnestic response while this will have no added value in our macaque infection model due to the relatively high challenge virus inoculum, and rapid clearance of the virus. In macaques, durable protection could be predicted based on humoral immune response levels. Passive transfer studies recently confirmed antibody responses as correlates of protection in rhesus macaques^18^. Whether binding and neutralizing antibody levels are a more universal correlate across multiple vaccine platforms and allow prediction of protection beyond 6 months is likely but will need to be confirmed. Ad26.COV2.S has recently shown an early indication of efficacy of 85% in humans with a median follow up of participants of 2 months^19^, confirming that the SARS-CoV-2 macaque model can be used to assess COVID-19 vaccine potency. Based on immunological follow-up, it will be ascertained whether predictions based on the macaque model are in line with observed protection in humans. If confirmed, durability of protection in humans could be estimated based on immunogenicity data. This may become especially relevant in the imminent situation where participants who were randomized to receive placebo in ongoing phase 3 efficacy trials will be offered vaccination, or when placebo recipients are lost to follow-up due to eligibility for vaccination in national vaccine campaigns and long term efficacy can no longer be evaluated in a blinded, placebo-controlled setting.

## Author Contributions

Designed studies and reviewed data: R.R, L.S., D.S., F.W., J.H., M.l.G., J.S., D.H.B., H.S. and R.Z. Performed experiments and analyzed data: J.S., R.S., S.K.R.H., J.E.M.v.d.L., L.D., D.N.C.C., N.G., S.J., S.T., A.C., N.M., J.Y. and W.K. Draft the paper: R.R., L.S., D.S. and W.K. Reviewed the paper: all authors.

## Acknowledgments

This project was funded in part by the Department of Health and Human Service Biomedical Advanced Research and Development Authority (BARDA) under contract HHS0100201700018C. We thank Gert Scheper, Martin Friedrich Ryser and Annemieke Manten for reviewing the paper and for providing valuable input. We thank the team of Bioqual (Rockville, MD, United States of America) for their work on the NHP durability study which involved challenged animals, in particular dr. Hanne Andersen. We thank the Nexelis Team of the Laval (Canada) and Seattle (United States of America) sites for their speed and flexibility in accommodating sample analyses within short timelines.

