## Supplementary Materials for "SARS-CoV-2 binding and neutralizing antibody levels after vaccination with Ad26.COV2.S predict durable protection in rhesus macaques"

### Supplementary figures and legends

#### Supplementary Figure 1: S-protein binding antibody levels from preceding vaccine studies

**a**

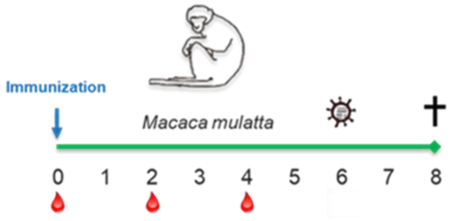


| Study | Immunization (week 0) | Construct | Dose (vp) | SARS-CoV-2 (week 6) | N |
|---|---|---|---|---|---|
| 20-07 | Ad26NCOV002 | tPA.S | 1x10^11 | $1x10^5$ $TCID_{50}$ USA-WA1/2020 | 4 |
| | Ad26NCOV004 | tPA.S.PP | 1x10^11 | | 4 |
| | Ad26NCOV006 | S | 1x10^11 | | 4 |
| | Ad26NCOV008 | S.dCT | 1x10^11 | | 4 |
| | Ad26NCOV014 | tPA.WT.S | 1x10^11 | | 4 |
| 20-07<br>20-09 | Ad26NCOV028 | S.dTM.PP | 1x10^11 | | 4<br>2 |
| 20-09<br>20-14 | Ad26.COV2.S (Ad26NCOV030) | S.PP | 1x10^11 | | 6<br>5 |
| 20-14 | | | 5x10^10 | | 5 |
| | | | 1.125x10^10 | | 5 |
| | | | 2x10^9 | | 5 |
| 20-07<br>20-09<br>20-14 | N/A | N/A | N/A | | 10<br>10<br>10 |

**b**

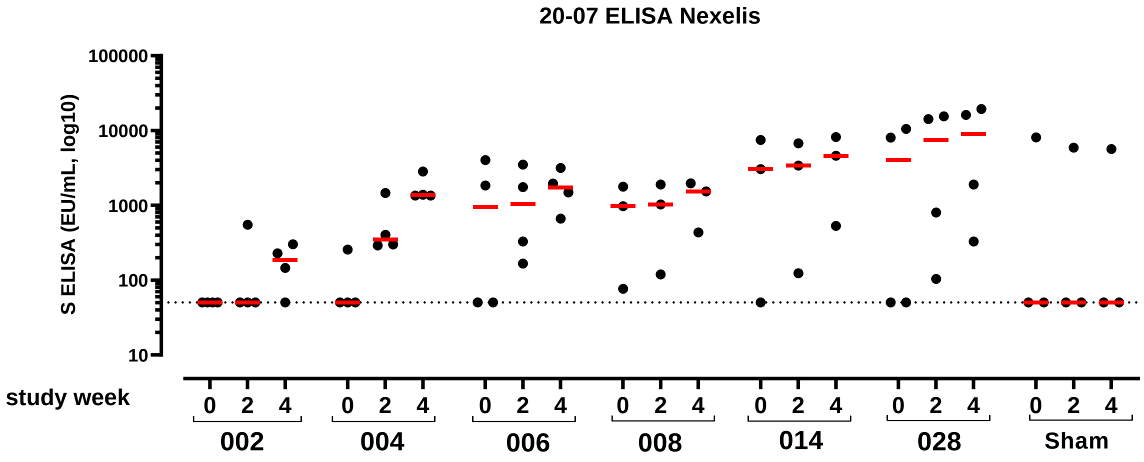

**c**

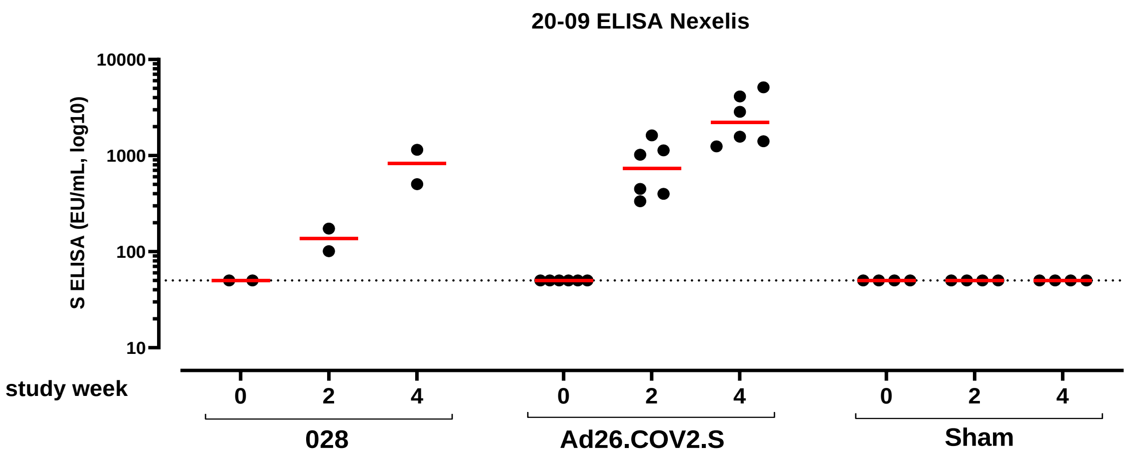


**d**

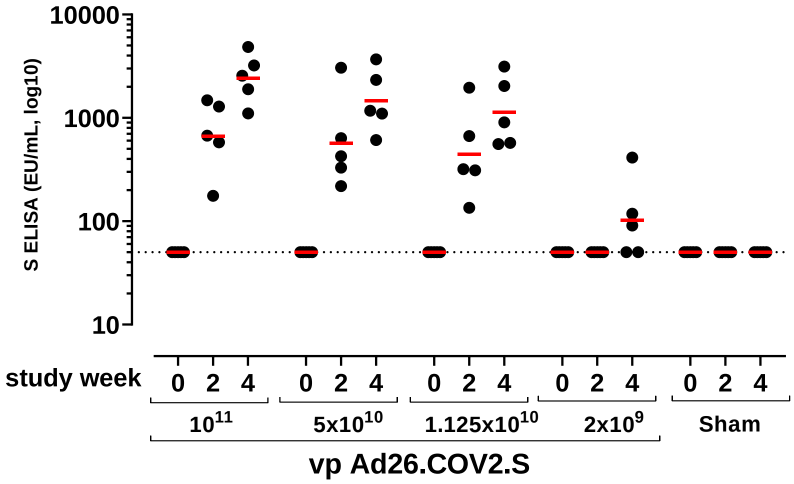


**Supplementary Figure 1: S-protein binding antibody levels from studies 20-07, 20-09, and 20-14. (a)** Schematic representation of the three studies. Groups of N monkeys received one-dose regimen of Ad26 based vaccine candidates. In total a group of 30 unvaccinated controls was included in the studies. Six weeks after the vaccination animals were challenged with $1x10^5$ TCID50 SARS-CoV-2 via the intranasal and intratracheal routes. Ad26 constructs: tPA.S (002) tissue plasminogen activator (tPA) leader sequence with full-length Spike protein (S), tPA.S.PP (004) tPA leader sequence with full-length S with mutation of the furin cleavage site and two proline stabilizing mutations, S (006) wildtype leader sequence with native full-length S, S.dCT (008) wildtype leader sequence with S with deletion of the cytoplasmic tail, tPA.WT.S (014) tandem tPA and wildtype leader sequences with full-length S, S.dTM.PP (028) wildtype leader sequence with S with deletion of the transmembrane region and cytoplasmic tail reflecting the soluble ectodomain, with mutation of the furin cleavage site, proline stabilizing mutations, and a foldon trimerization domain, S.PP (030 / Ad26.COV2.S) wildtype leader sequence with full-length S with mutation of the furin cleavage site and proline stabilizing mutations. **(b, c, d)** S-protein binding antibody levels (S-ELISA, EU/ml, log10) were determined at baseline, 2-, 4- and 6-weeks post-vaccination. Some animals in the candidate selection study (20-07) had background reactivity prior to vaccination (week 0). Background reactivity was not observed in psVNA, and likely represents aspecific binding. Assay LLOQ is shown as dashed lines. Horizontal bars indicate geometric mean of response within each group.

**Supplementary Figure 2: Correlation of cellular immune responses with protection against SARS-CoV-2 in the lung and the nose.**

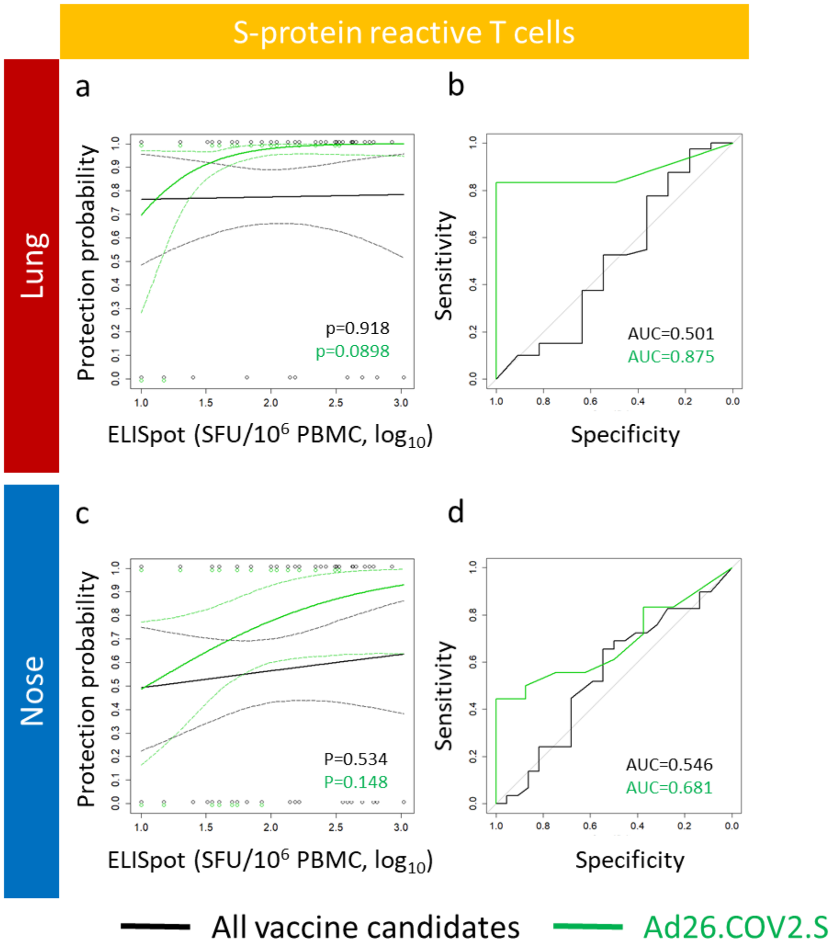


(**a, c**) Logistic models of the correlation between the level of S-protein specific T cells (IFN-gamma ELISpot, SFU/$10^6$ PBMC, $\log_{10}$) and protection against viral load in lung (a, BAL) and nose (c, swabs), based on the dataset of all Ad26-based vaccine candidates combined (black line) and Ad26.COV2.S alone (green line). 95% confidence intervals are represented by dashed lines in the same color. Individual datapoints (y=0: detectable viral load; y=1 undetectable viral load) are represented by open circles in the same color. (**b, d**) ROC curves for the logistic models presented in panels a and c, respectively. Area under the ROC curve (AUC) is indicated and represents a measure of the sensitivity and specificity of the logistic model.

**Supplementary Figure 3: Viral load kinetics and duration in the lungs of individual animals of the 6M durability study**

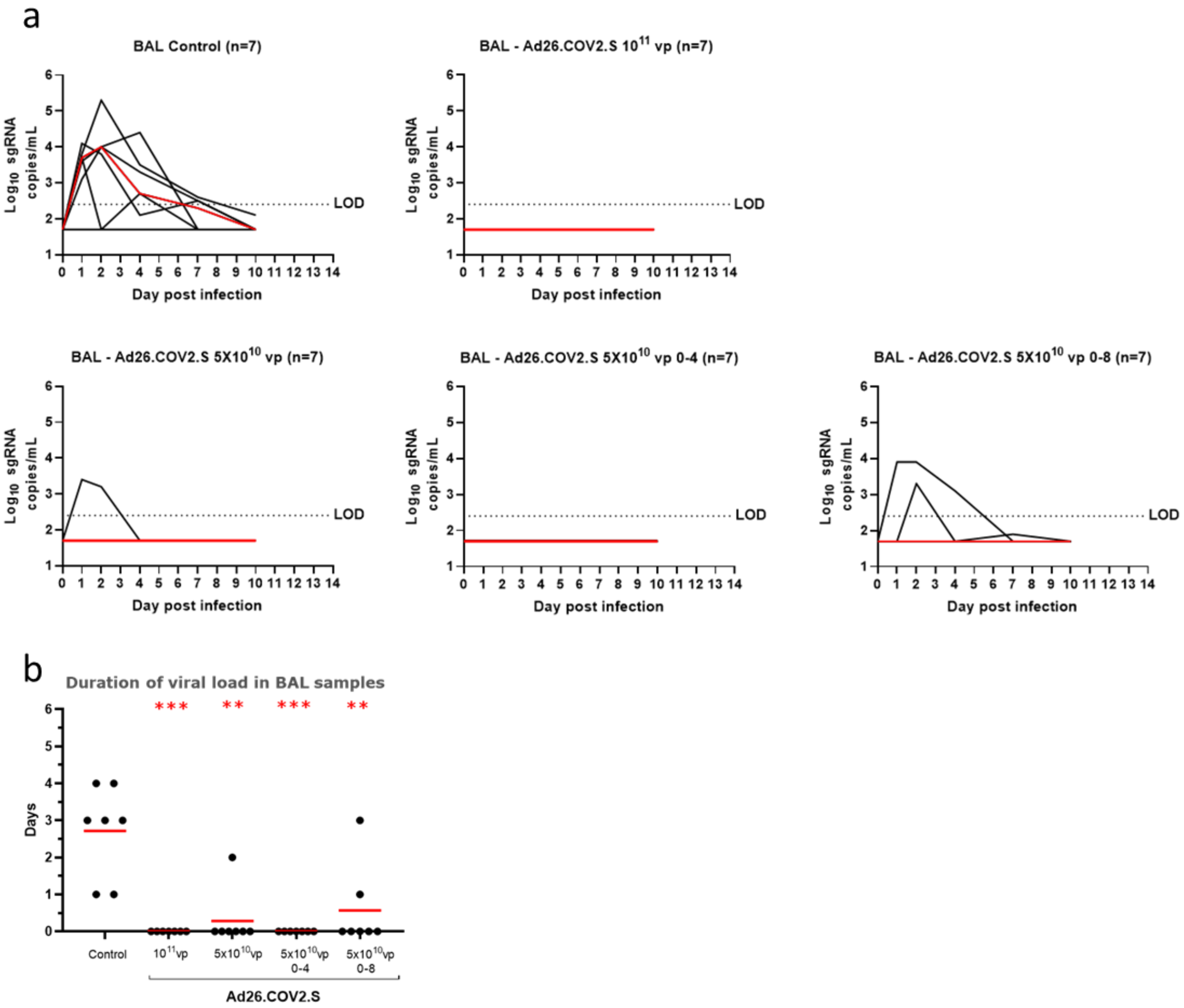


**(a)** Viral load (sgmRNA copies/mL) kinetics in BAL samples after SARS-CoV-2 challenged of control and vaccinated rhesus macaques. Black lines represent individual animals, red lines the group median. Assay LOD is shown as a dashed line. **(b)** Duration (days) of viral load in BAL samples after SARS-CoV-2 challenged of control and vaccinated macaques. Black dots represent individual animals, red lines the group mean. Red asterisks indicate the statistical significance. (*p<0.05, ** p<0.01, *** p<0.001). The statistical analysis was performed using the Mann-Whitney U test.

#### Supplementary Figure 4: Viral load kinetics in the nose of individual animals of the 6M durability study

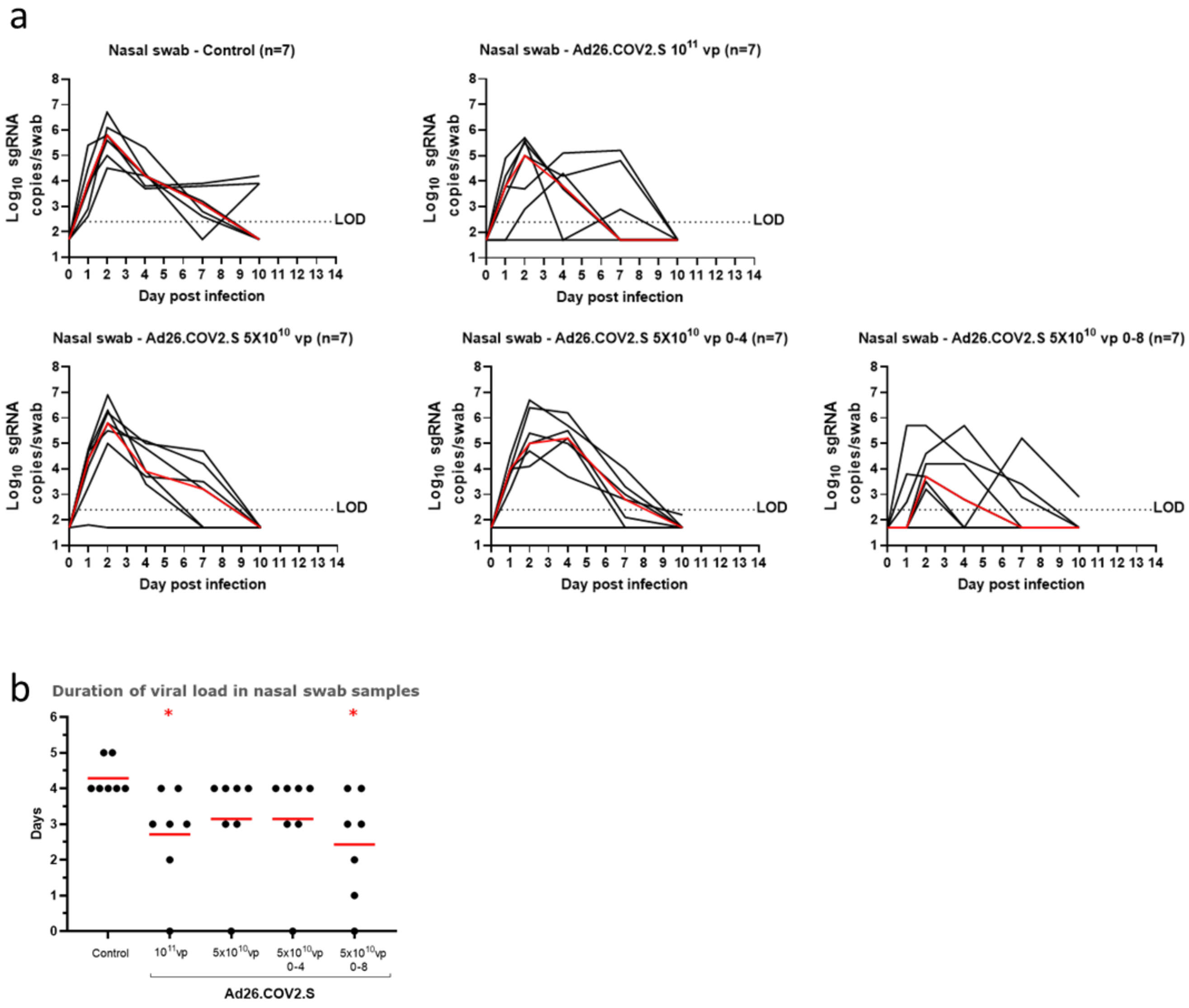


**(a)** Viral load (sgRNA copies/swab) kinetics in nasal swab samples after SARS-CoV-2 challenged of control and vaccinated rhesus macaques. Black lines represent individual animals, red lines the group median. Assay LOD is shown as a dashed line. **(b)** Duration (days) of viral load in nasal swab samples after SARS-CoV-2 challenged of control and vaccinated macaques. Black dots represent individual animals, red lines the group mean. Red asterisks indicate the statistical significance. (*$p<0.05$, ** $p<0.01$, *** $p<0.001$). The statistical analysis was performed using the Mann-Whitney U test.

**Supplementary Figure 5: Comparison between observed and predicted protection based on correlate models obtained with data of all vaccine candidates combined.**

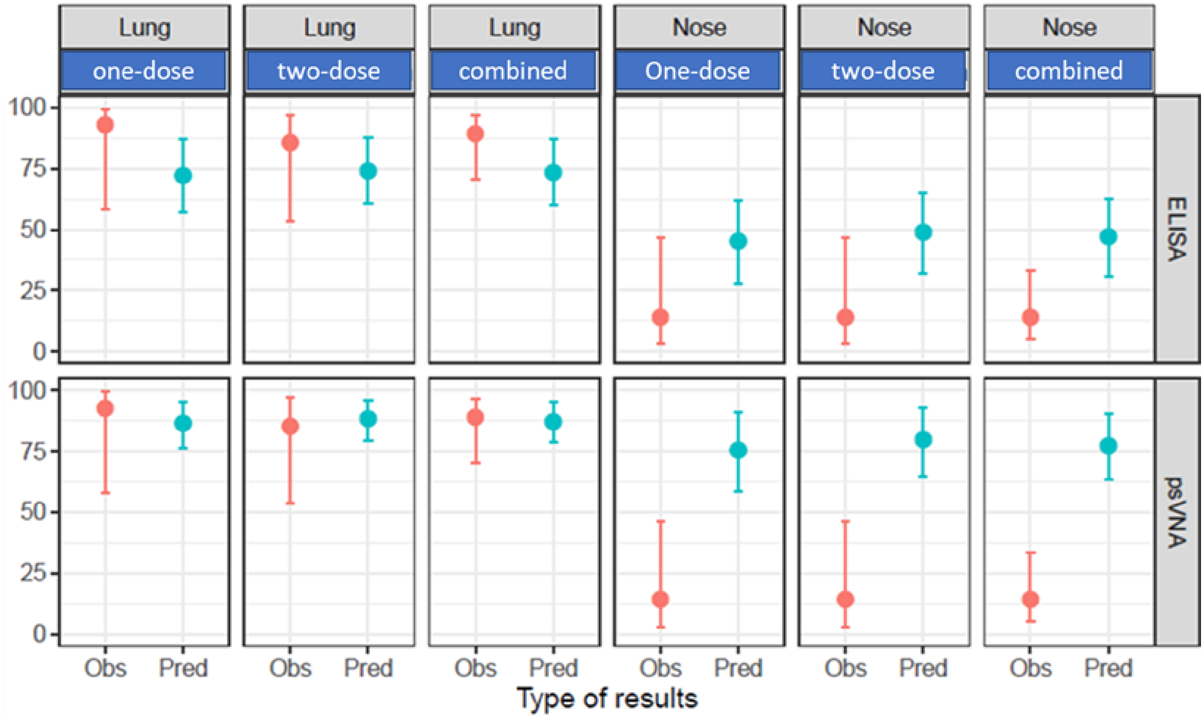


Comparison of the observed (Obs) protection probability 6 months after vaccination with Ad26.COV2.S with the predicted (Pred) protection probability based on pre-challenge binding (S-ELISA) and neutralizing antibody (psVNA) levels and correlate of protection models (logistic models) constructed based on immunogenicity data of all Ad26-based vaccine candidates, obtained 4 weeks after vaccination. Dots are the mean values and the vertical lines represent the 95% CI. Predictions are shown for logistic models based on data of all vaccine candidates in both the lung and the nose. Data from the durability study were grouped by one-dose regimes (one-dose, n=14), two-dose regimens (one-dose, n=14), or analyzed together (combined, n=28).

**Supplemental methods:**

***Statistical Methods***

Peak and duration of viral load in BAL and nasal swabs samples of each vaccinated group were compared to the control group using Mann-Whitney U test.

Protection outcome is binary and driven by the viral load observation (ie, defined as 0 if the macaque had a detectable viral load as measured by sgRNA and as 1 if the macaque had an undetectable viral load). The effect of the different immunogenicity response markers on the measure of protection is investigated using a logistic regression modeling approach. Penalized logistic models were built using Firth's method,[1] with protection outcome as the dependent variable and immune response level as the independent variable. A bootstrap procedure was applied where the dataset was resampled 10,000 times with replacement and the logistic model was fitted to each resampled dataset. The 2.5th and 97.5th percentiles of the fitted protection probabilities were calculated over a grid of immune response values to generate the pointwise 95% confidence band.

The above developed logistic model can be used to predict the probability of protection for a set of newly measured immunogenicity samples for which we want to have an idea about their probability of protection without having the actually observed protection status, an example of such a set of new samples are the pre-challenge immunogenicity results for the vaccinated animals. This was done using a double bootstrap procedure. In this procedure both the macaque dataset used to fit the logistic model as well as the dataset with new samples of interest are resampled 10,000 times each with replacement. The logistic regression model describing the effect of the immunogenicity marker on the measure of protection, is refitted to each resampled macaque dataset and used to predict the probability of protection for the resampled dataset with new samples. As a result, 10,000 mean predicted protection probabilities for the resampled dataset with new samples is obtained. The 95% CI is then derived by taking the $2.5^{th}$ and $97.5^{th}$ percentiles of the 10,000 mean predicted protection probabilities.

The estimated mean population probability of protection for the challenged macaques, together with a 95%CI based on their viral load measurements after challenge, are analyzed using a logistic regression model with overall intercept only using the glm functionality (lme4 package R),[2] and compared to the observed proportion of protection.

Simulations reported here were conducted using R version 3.6.0[3] on an x86_64-redhat-linux-gnu (64-bit) platform running under a Red Hat Enterprise Linux Server 7.4 (Maipo) using Rstudio Version 1.1.453.[4] The seed was set to allow reproducibility.

***SARS-CoV-2 Pseudotyped Virus Neutralization Assay on Vero E6 Cells***

The SARS-CoV-2 pseudoviruses expressing a luciferase reporter gene are generated in an approach similar to as described previously.[5-7] Briefly, the packaging construct psPAX2, luciferase reporter plasmid pLenti-CMV Puro-Luc and spike protein expressing pcDNA3.1-SARS CoV-2 SΔCT are co-transfected into HEK293T cells. The supernatants containing the pseudotype viruses are collected and stored at -80°C until use. To determine the neutralization activity of the antisera from vaccinated animals, HEK293T-hACE2 stable cells are seeded in 96-well tissue culture plates. Serial dilutions of serum samples are prepared and mixed with pseudovirus. The mixture is incubated at 37°C for 1 hour before adding to HEK293T-hACE2 cells. 48 hours after infection, cells are lysed in Steady-Glo Luciferase Assay according to the manufacturer's instructions. SARS-CoV-2 neutralization titers are defined as the sample dilution at which a 50% reduction in RLU is observed relative to the average of the virus control wells. Assay LOD=20 IC50 (1.3 log10). This assay was previously shown to strongly correlate with wtVNA, and SARS-CoV-2 Spike IgG ELISA.[8]

***SARS-CoV-2 Spike IgG ELISA***

SARS-CoV-2 Spike-specific binding antibody concentrations was determined using the human SARS-CoV-2 Spike immunoglobulin G (IgG) ELISA. The SARS-CoV-2 antigen used is a stabilized pre-fusion spike protein ((2P), Δfurin, T4 foldon, His-Tag) produced in ES-293 cells. The ELISA was performed at Nexelis (Seattle, WA, USA).

In brief, The SARS-CoV-2 Pre-Spike IgG ELISA is an indirect ELISA which is based on the antibody/antigen interactions. Purified SARS-CoV-2 Pre-Spike Antigen is adsorbed to the wells of a microplate. Diluted serum samples (test samples, standard, and quality controls) are added in the wells. Anti-SARS-CoV-2 Pre-Spike antibodies if present in the serum samples bind to the immobilized SARS-CoV-2 Pre-Spike antigen. Unbound sample is then washed from the wells, and enzyme-conjugated anti-human IgG is added. The anti-human IgG enzyme conjugate binds to the antigen-antibody complex. Excess conjugate is washed away, and 3,3′,5,5′-Tetramethylbenzidine (TMB) colorimetric substrate is added. Bound enzyme catalyzes a hydrolytic reaction, which causes color development. After the established time period, the reaction is stopped. The intensity of the generated color is proportional to the amount of anti-SARS-CoV-2 Pre-Spike antibodies bound to the wells. The optical density results are read on a spectrophotometer (ELISA plate reader). A reference standard on each tested plate is used to quantify the amount of antibodies against SARS-CoV-2 Pre-Spike in the sample according to the unit assigned by the standard (ELISA Laboratory Unit per milliliter: ELU/mL). Assay LOD is 3.4 EU/ml (0.53 $\log_{10}$ EU/ml) and LLOQ is 50.3 EU/ml (1.7 $\log_{10}$ EU/ml). This assay was previously shown to strongly correlate with wtVNA, and psVNA.[8]

***Viral Load Determination***

The presence of virus in BAL and nasal swab samples is measured by real-time polymerase chain reaction (RT-PCR) of SARS-CoV-2 E gene subgenomic ribonucleic acid (sgmRNA). The assay (based on Wölfel et al, 2020[13]) thus detects replicating virus and largely distinguishes between virus present in the inoculum, and infected cells. A standard curve is generated using RNA transcribed from the SARS-CoV-2 E gene sgmRNA cloned into a pcDNA3.1 expression plasmid. Prior to RT-PCR, samples are thawed at room temperature or 4˚C and reverse-transcribed using Superscript III VILO (Invitrogen) according to the manufacturer's instructions. The complementary deoxyribonucleic acid (cDNA) is stored at 4˚C until use. A Taqman custom gene expression assay (ThermoFisher Scientific) was designed using the sequences targeting the E gene sgmRNA. The PCR is carried out on a QuantStudio 6 and 7 Flex Real-Time PCR System (Applied Biosystems) according to the manufacturer's specifications. Standard curves were used to calculate sgRNA copies and were multiplied by 250 to get to copies per mL or per swab. Quantitative sensitivity (LOQ) per PCR reaction was determined as one copy, corresponding to a limit of detection (LOD) of 250 copies per sample.

3 R: A language and environment for statistical computing (R Foundation for Statistical Computing, Vienna, Austria., 2019).

4 RStudio: Integrated Development for R (RStudio, Inc., Boston, MA, 2016).

5 Chandrashekar, A. *et al.* SARS-CoV-2 infection protects against rechallenge in rhesus macaques. *Science* **369**, 812-817, doi:10.1126/science.abc4776 (2020).

6 Yu, J. *et al.* DNA vaccine protection against SARS-CoV-2 in rhesus macaques. *Science* **369**, 806-811, doi:10.1126/science.abc6284 (2020).

7 Yang, Z. Y. *et al.* A DNA vaccine induces SARS coronavirus neutralization and protective immunity in mice. *Nature* **428**, 561-564, doi:10.1038/nature02463 (2004).

8 Solforosi, L. *et al.* Immunogenicity and protective efficacy of one- and two-dose regimens of the Ad26.COV2.S COVID-19 vaccine candidate in adult and aged rhesus macaques. *bioRxiv*, 2020.2011.2017.368258, doi:10.1101/2020.11.17.368258 (2021).
